# Questioning the role of selected somatic *PIK3C2B* mutations in squamous non-small cell lung cancer oncogenesis

**DOI:** 10.1101/164749

**Authors:** Marcus Kind, Jolanta Klukowska-Rötzler, Sabina Berezowska, Alexandre Arcaro, Roch-Philippe Charles

**Affiliations:** University Children’s Hospital Bern, Freiburgstrasse 31, 3010 Bern, Switzerland; Department of Emergency Medicine, University Hospital Bern, Freiburgstrasse 16c,3010 Bern, Switzerland; Institute of Pathology, University of Bern, Murtenstrasse 31, 3008 Bern,Switzerland; Institute of Biochemistry and Molecular Medicine, and Swiss National Center of Competence in Research (NCCR)TransCure, University of Bern, Bühlstrasse 28, 3012 Bern, Switzerland

**Keywords:** PI3K, *PIK3CA*, *PIK3C2B*, lung cancer, Sanger sequencing

## Abstract

PI3K signaling is frequently dysregulated in NSCLC-SQCC. In contrast to well characterized components of the PI3K signaling network contributing to the formation of SQCC, potential oncogenic effects of alterations in *PIK3C2B* are poorly understood. Here, a large cohort (n=362) of NSCLC-SQCC was selectively screened for four reported somatic mutations in *PIK3C2B* via Sanger sequencing. In addition, two mutations leading to an amino acid exchange in the kinase domain (C1181, H1208R) were examined on a functional level for their oncogenic potential.

None of the mutations were identified in the cohort while well characterized hotspot *PIK3CA* mutations were observed at the expected frequency. Ultimately, kinase domain mutations in PI3KC2β were found to have no altering effect on downstream signaling. A set of SQCC tumors sequenced by The Cancer Genome Atlas (TCGA) equally indicates a lack of oncogenic potential of the kinase domain mutations or *PIK3C2B* in general.

Taken together, this study suggests that *PIK3C2B* might only have a minor role in SQCC oncogenesis.

## Introduction

Phosphoinositide-3-kinases (PI3Ks) are able to phosphorylate the inositol ring of three different phosphatidylinositol lipid substrates (PtdIns, PtdIns4P, PtdIns(4,5)P_2_), minor compounds on the cytosolic site of eukaryotic cell membranes. Following activation by upstream agonists such as receptor tyrosine kinases (RTKs) or G protein coupled receptors (GPCR), PI3Ks generate 3-phosphoinositides as second messengers. These 3-phosphinositides coordinate the function and localization of numerous effector proteins. Downstream pathways of those proteins control a broad range of different physiological functions as diverse as proliferation, migration, apoptosis and cell metabolism (1–5). Eight different catalytic PI3K isoforms have been described that are subdivided into three different classes (class I, class II and class III). This classification is based on substrate specificity, associated co-factors and sequence homologies.

Because of its central role in intracellular signaling, dysregulation of the PI3K network belongs to the most common events in human cancers (6). Prominent examples are loss of function mutations in PTEN, the main PI3K phosphatase and antagonist of PI3K signaling (7,8). In regard to PI3K isoforms, there is plenty of evidence that alterations in class I alpha (p110_α_) promote oncogenesis. Somatic mutations clustered in *PIK3CA* hotspot regions are frequently found in a wide array of cancers (9,10) and their oncogenic potential is well documented in functional studies (11,12). Aside from p110_α_, there are numerous publications linking further PI3K isoforms to tumorigenesis (13–15).

Among them is the class II isoform C2β. As class II PI3Ks were discovered based on sequence homologies with class I and class III instead of a functional context, the physiological role and downstream pathways of PI3KC2β remain enigmatic. Nevertheless, PI3KC2β has been repeatedly associated with various steps of oncogenesis in different cell lines. These range from enhanced cell migration (16) to an increase in chemotherapeutic resistance (17), anchorage-independent growth (18) and cell proliferation (19). Moreover, a study characterizing the exomes of 31 non-small cell lung cancer (NSCLC) genomes found 4 missense mutations in *PIK3C2B*: c.349C>G (P117A), c.3542G>T (C1181F), c.3623A>G (H1208R) and c.4407G>T (L1469F). Two of them were located in a highly conserved region of the kinase domain (C1181 and H1208R) (20). The frequency was even higher (3/12) when only considering the squamous cell carcinoma (SQCC) subtype.

Together with adenocarcinomas (ADC), SQCC comprise the majority of all non-small cell lung carcinomas (NSCLC) (21). Recent efforts have been undertaken to unveil the underlying changes in the genome, transcriptome and proteome of these two histological subtypes. This led to growing evidence of distinct genomic alteration patterns. As for SQCC, oncogenesis appears to rely on alterations in squamous differentiation (22), oxidative stress response (23) and PI3K signaling (24). According to a large genomic analysis, aberrant PI3K signaling is present in approximately half of all cases (25), mainly through alterations in *PIK3CA* and *PTEN*.

Given the high prevalence of *PIK3C2B* mutations in the small NSCLC cohort screened by Liu *et.al.* and the connection to multiple steps of cancer progression, *PIK3C2B* and the reported kinase domain mutations C1181F and H1208R were closer examined in regard to promote aberrant PI3K signaling in NSCLC-SQCC.

For this purpose, a cohort of 362 NSCLC-SQCC was selectively screened for all four reported alterations in *PIK3C2B.* To embed the sequencing results into a broader context, clinical outcomes of a set of SQCC sequenced by the cancer genome atlas (TCGA) (26,27) were analyzed with respect to alterations in *PIK3C2B*. Moreover, the functional impact of C1181F and H1208R was assessed in relation to its potential to hyper-activate downstream PI3K signaling in HEK293 cells.

## Material and Methods

### DNA isolation from tumor samples

Punches from paraffin embedded NSCLC-SQCC tumor samples were provided by the Institute of Pathology and Tumor Tissue Bank, University of Bern. The SQCC cohort included 362 primary resected tumors and 29 corresponding mediastinal lymph node metastases diagnosed at the Institute of Pathology 2000-2013. In order to exclude pulmonary metastases of other SQCC, patients with previous SQCC of other organs were not included. The cohort comprised 52 females and 310 males with a median age of 69 years at the time of operation (range 43-85 years of age) and included all UICC 2009 pT stages (pT1a=34, pT1b=49, pT2a=119, pT2b=53, pT3=77, pT4=30) and UICC 2009 tumor stages (IA=61, IB=79, IIA=73, IIB=51, IIIA=81, IIIB=8, IV=8). The study was approved by the Cantonal Ethics Commission of the Canton of Bern (KEK200/14), which waived the requirement for written informed consent. DNA was isolated from one or two paraffin punches per sample by using Qiaamp DNA MicroKIT kits (Qiagen, cat. no. 56304).

### Sanger sequencing

Potentially mutated sites were amplified via AmpliTaq Gold DNA Polymerase (ThermoFischer, cat. no. N8080241) in a thermocycler (UNO II, Biometra). Cycling conditions consisted of an initial denaturation step at 95°C for 10 min and 30 cycles of denaturation (95°C, 30 sec), annealing (60°C, 30 sec) and extension (72°C, 40 sec). Primer sequences and amplification conditions for *PIK3CA* screening were adopted from Samuels *et. al.* (10).

Following amplification of the regions of interest, 5phosphates of the PCR products were degraded with rAPid alkaline phosphatase (Roche, cat. no. 4898133001) followed by 25 cycles of forward or reverse amplification at the same cycling conditions as indicated above (BigDye® Terminator v3.1 Cycle Sequencing Kit, Life Technologies, cat. no. 4337455). After DNA precipitation, amplicons were dissolved in Hi-Di formamide (Thermo-Scientific, cat. no. 4311320) and sequenced with an ABI3730 DNA analyzer (Applied Biosystems).

### Primers

Primers of this project (Table1) were purchased from Microsynth and designed with the Primer-Blast web tool (ncbi. nlm.nih.gov/tools/primer-blast/).

**Table1:**
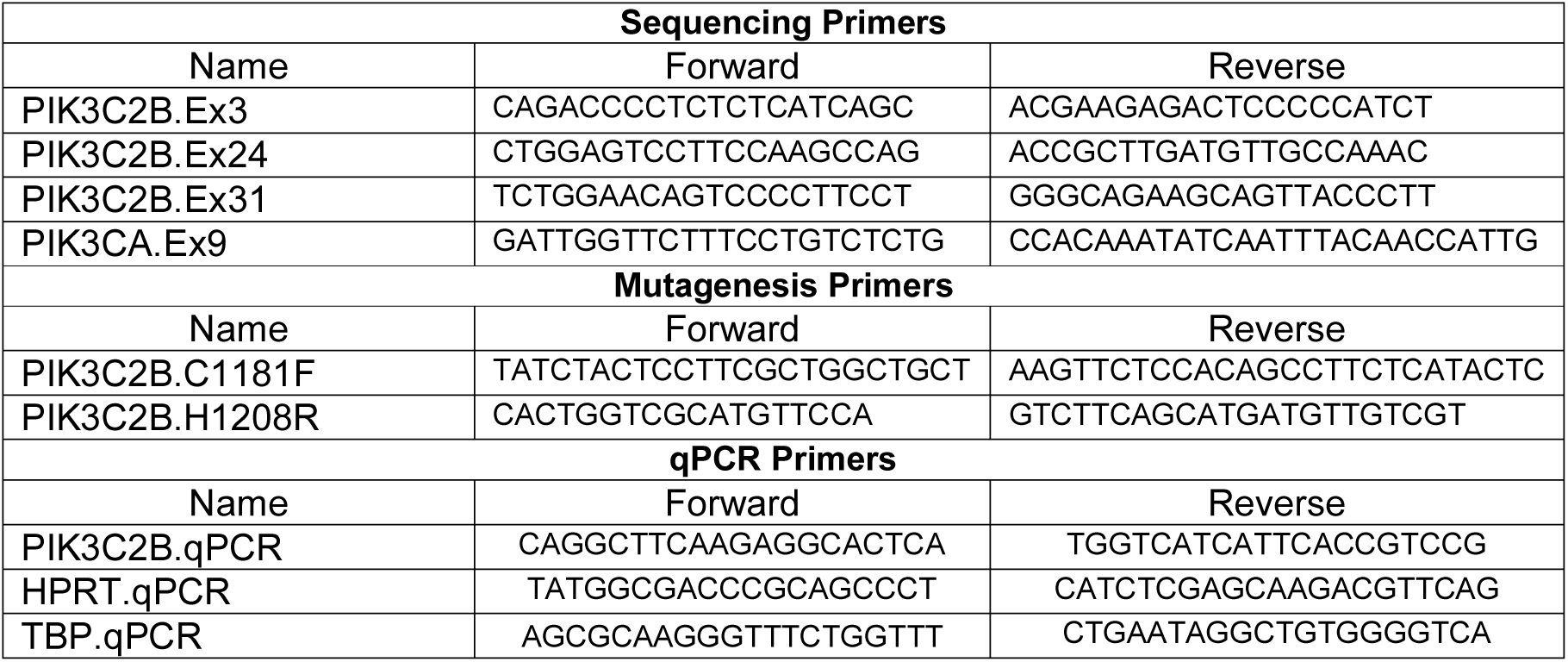
Primers used for sequencing and quantification.

### Plasmid engineering

A *PIK3C2B* expression vector with a C-terminal Myc-Tag was purchased from Origene (cat. no. NM-002646). Primers with an adequate nucleotide mismatch were designed to engineer C1181F and H1208R amino acid exchanges into the plasmid. The implemented changes in the base triplicates were: C1181F: TGC>TTC / H1208R: CAC>CGC. To facilitate ensuing ligation, primers were additionally phosphorylated at the 5`end.

Site-directed mutagenesis was carried out with the Phusion Site-Directed Mutagenesis kit (Thermo Scientific, cat. no. F541). Mutations were incorporated by following the indicated cycling conditions: initial denaturation (98°C, 10 min) was followed by 25 cycles of annealing (C1181F: 69°C, H1208R: 64,5°C, 20 sec) and extension (72°C, 5 min). PCR products were ligated (Promega, cat. no. M180S) and successful engineering was tested via Sanger sequencing.

An empty control plasmid was created by removing the *PI3KC2B* open reading frame via restriction digestion with NheI (Promega, cat. no. R650A) and MluI (Promega, cat. no. R638A). Plasmid fragments were separated in 1 % agarose gel and purified (Promega, cat. no. A9281). Afterwards, 5`overhangs were blunted (NEB, cat. no. M0210S) and ligated (Promega, cat. no. M180S). Plasmid constructs were cloned into E. Coli XL-1 Blue bacteria.

### Bacterial transformation

Competent E.Coli XL-1 Blue bacteria were transformed with 150 ng of target plasmid by applying a 42°C heat shock for 85 sec. After overnight culture in LB medium containing adequate antibiotic concentration (100 μg/ml ampicillin), clonally expanded plasmids were isolated with Pureyield^TM^ Plasmid Miniprep/ Midiprep kits (Promega, cat. no. A1223/A2495).

### Cell lines and culture

HEK293 cells were purchased from the American Type Culture Collection (ATCC). Cells were cultured in Dulbecco’s Modified Eagle Medium (Sigma Aldrich, cat. no. D5796) supplemented with 10 % FBS (Gibco, cat. no. 10082147), 2 mM L-glutamine (Gibco, cat. no. 25030081) and 50.000 units of penicillin/streptomycin (Gibco, cat. no. 15140122). Cells were kept up to passage 50 or 3 months maximum.

### Transient transfection

HEK293 cells were transfected at 50-60 % confluency in different formats (6 well / 10 cm) with calcium phosphate. Appropriate amounts of plasmid DNA (6 well: 4 μg / 10 cm dishes: 30 μg) were thoroughly mixed with 1/10 Vol. of 2.5 M CaCl_2_ and 2x HEPES buffered saline (HBS, 40 mM HEPES, 10 mM D-Glucose, 10 mM KCl, 270 mM NaCl, 1,5 mM Na_2_HPO_4_). Subsequently, the transfection mix was added dropwise to HEK293 cells. After overnight exposure to the precipitate, medium was changed and cells were further cultivated for 48-72 h.

### qPCR

RNA from transfected HEK293 cells was isolated with the RNeasy Mini kit (Qiagen, cat. no. 74104), followed by reverse transcription (Applied Biosystems, cat. no. 4368814). Quantitative PCRs were performed in a ViiA7 cycler (Applied Biosciences) using SybrSelect Mastermix (Applied Biosystems, cat. no. 4472908).

Expression of mRNA was normalized to TATA box binding protein (*TBP)* and hypoxanthine-guanine phosphoribosyl transferase (*HPRT)* housekeeping genes.

### Western blot

Proteins were extracted in RIPA buffer (20 mM Tris-base pH 8, 150 mM NaCl, 1 % Triton-X-100, 0.1 % SDS, 0.5 % sodium deoxycholate) supplemented with 100 µM Na_3_VO_4_, 25 mM β-glycerophosphate, 1 mM NaF and cOmplete^TM^ Protease Inhibitor Cocktail (Roche, cat. no. 11836170001). Pierce BCA protein assay kit (Thermo Scientific, cat. no. 23225) was used to determine protein concentration. Subsequently, 20 µg of protein were separated via SDS-PAGE, transferred onto nitrocellulose membranes and blocked in Tris buffered saline (TBS, 130 mM NaCl, 30 mM Tris-Cl, pH 7.5) containing 5 % Bovine Serum Albumin (BSA) for 2 h. Western blots were probed with rabbit anti-PI3KC2β polyclonal antibody (1/1000, described in (28)) rabbit anti-P-Akt Ser473 (1:1500, Cell signaling technology, cat. no. 4060L), rabbit anti-P-S6 Ser240/244 (1:2500, Cell signaling technology, cat. no. 5364L), mouse anti-total-Akt (1:2000, Cell signaling technology, cat. no. 2920S) mouse anti-total-S6 (1:2000, Cell signaling technology, cat. no. 2317S) and mouse anti-β-actin antibody (1/2000, Sigma Aldrich, cat. no. A5316). Primary antibodies were detected by using goat anti-rabbit IR680 (1/10.000, cat. no. 926-68071, LiCor Bioscience) and goat anti-mouse IR800 (1/10.000, cat. no. 926-32210, LiCor Bioscience) antibodies and analyzed with a LI-COR OdysseySa imaging system.

### Immunoprecipitation and lipid kinase assay

Lipid kinase activity of exogenously expressed PI3KC2β was measured with a bioluminescence based kit purchased from Promega (ADP-Glo-Kinase Assay, cat. no. V6930).

HEK293 cells were grown in 10 cm diameter dishes and transfected with plasmid constructs as described above. Cells were lysed for 20 minutes on ice by applying 2 ml lysis buffer (1 % Triton X-100, TrisCl 50 mM pH 7.4, NaCl 150 nM, 1 mM EDTA), supplemented with 100 μM Na_3_VO_4_, 1 mM NaF, 20 mM β- glycerophosphate and cOmplete^TM^ Protease Inhibitor Cocktail. Then, lysates were centrifuged (16’000 g, 4°C, 30 min) to remove insoluble cellular debris and supernatant was incubated with anti-MycTag antibodies for 2 hours at 4°C under continuous agitation. Antibody-protein complexes were pooled down by incubation (1 h, 4°C), with sepharose beads (GE Healthcare, cat. no. 17061801) under continuous agitation. Immunoprecipitates were washed 3 times in lysis buffer, followed by quick spin down (4000 g, 4°C, 1 min). Ensuing, sepharose pellets were re-suspended in kinase reaction buffer (40 mM Tris HCl pH 7.5, 20 mM MgCl_2_, 0.1 mg/ml BSA), supplemented with 0.2 mg/ml phosphatidylinositol substrate (PI, Sigma Aldrich, cat. no. 79403) and incubated on ice for 20 min. Enzymatic reaction was started after addition of 50 μM ATP and precipitates were incubated for 30 min at room temperature. Remaining experimental steps were carried out according to the manufacturer’s protocol. Luminescence was measured with a Modulus Microplate reader (Turner Biosystems).

### Immunofluorescence

HEK293 cells were grown on glass coverslips. After 10 % formalin fixation (10 min), coverslips were washed 3x10 min in phosphate buffered saline (1x PBS: 137 mM NaCl, 2.7 mM KCl, 18 mM KH_2_PO_4,_ 10 mM Na_2_HPO_4_) and cells were subsequently permeabilized with a 1x PBS, 0.3 % Triton-X100 solution. Following blocking with a 1 % BSA, 0.2 % gelatin, 0.05 % saponin in 1x PBS solution and washing with a 0.1 % BSA, 0.2 % gelatin, 0.05 % saponin in 1x PBS solution, fixed cells were treated overnight at 4°C with a mouse anti-MycTag (9E10 epitope) antibody diluted in an adequate buffer (0.1 % BSA, 0.1 % sodium azide, 0.3 % triton X-100 in 1x PBS). Then, coverslips were rinsed 3 times in washing solution. Cells were further incubated with a fluorescent secondary anti-mouse-Alexa647 antibody (1:500, ThermoFischer, cat. no. A32728) to detect antigen-antibody complexes and counter-stained with DAPI (500 ng/ml, Sigma Aldrich, cat. no. 32670-25MG). Slides were scanned with a Pannoramic Midi II scanner (3DHISTECH Ltd.).

### In silico meta-analysis

A SQCC data set sequenced by the TCGA and available on cbioportal (26,27) was analyzed in relation to *PIK3C2B* sequencing information. Raw data were visualized and formatted with Excel and Prism7 (GraphPad).

### Statistical analyses

Statistical analyses were conducted using Prism 7. The statistical test used is indicated in the legend of the figure. A value of p<0.05 was considered to be significant.

## Results

### Cohort validation

To ensure robust sequencing data, it had to be ascertained, that assay sensitivity was sufficient to detect tumor specific mutations and that the obtained cohort was representative. To validate the former, different ratios of *PIK3C2B*^*WT*^ and *PIK3C2B*^*A^3623^G*^ plasmids were analyzed via Sanger sequencing. Detection of 10 % mutated plasmid combined with 90 % wildtype plasmid was possible (Fig. 1B).This result was considered to be sufficient for further analyses, as paraffin punches contained tumor fractions > 30 %. To ensure that the cohort was representative, it was screened for the charge reversing hotspot mutations p110α^E542K^ and p110α^E545K^. Both mutations were present in the cohort (Fig. 1C & 1D). Relying on mutation data from the COSMIC database (cancer.sanger.ac.uk), a subsequent χ^2^ test revealed no significant difference between the observed and expected frequencies in the screened cohort (E542K p=0.383, E545K p=0.475).

**Figure 1:**
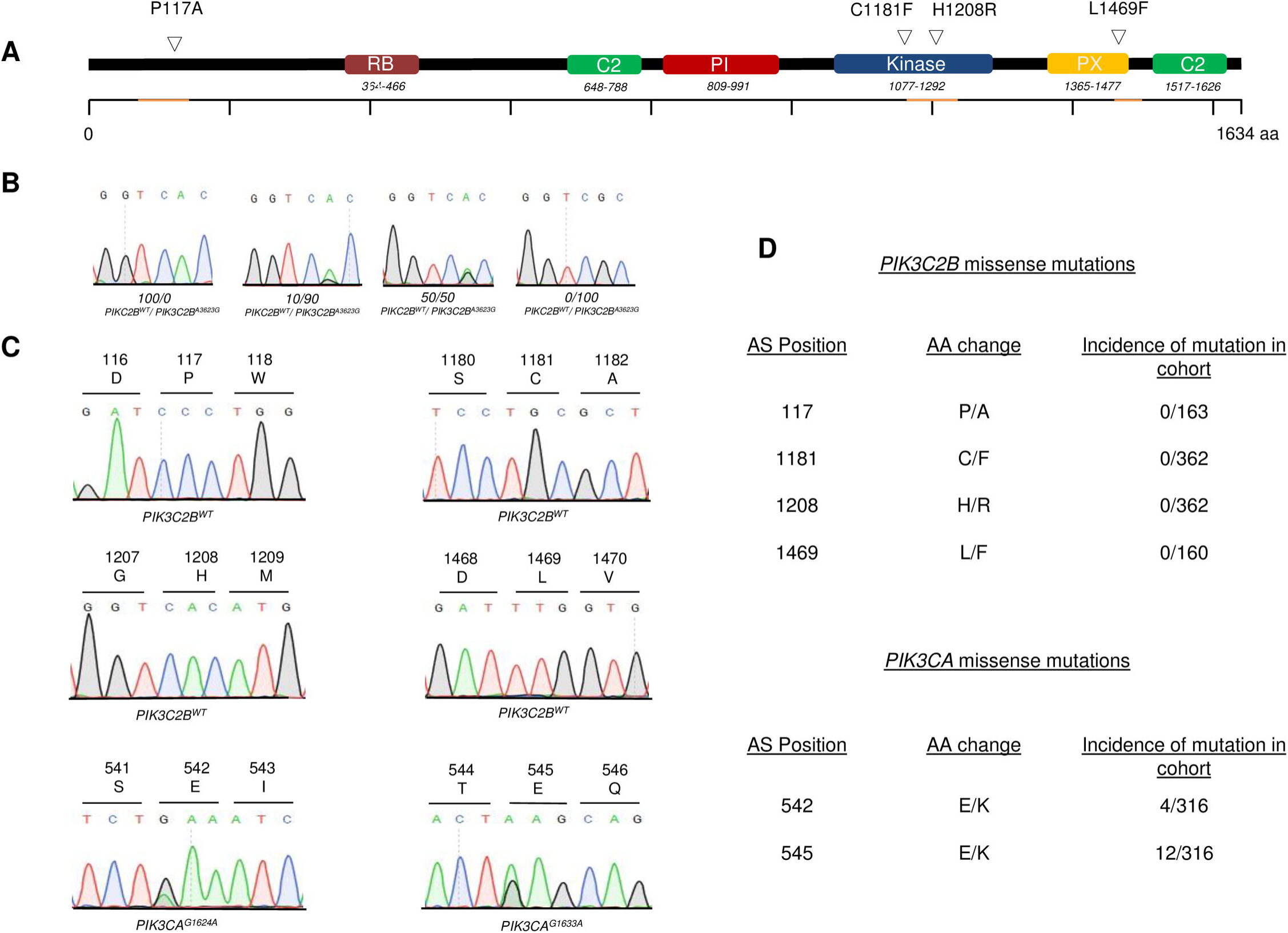
Screening of NSCLC-SQCC tumors for somatic *PIK3C2B* mutations. **A** Structural domains of *PIK3C2B* with reported mutations and sequenced regions (highlighted in orange). **B** Chromatograms of different *PIK3C2B*^*WT*^ */ PIK3C2B*^*^A3623G^*^ *ratios*. **C** Representative chromatograms of screened genomic regions in *PIK3C2B* and *PIK3CA*. **D** Table with incidence and position of found alterations in *PIK3C2B* and *PIK3CA*.

### PIK3C2B screening

After isolation of DNA from paraffin embedded tissue, samples were screened for the reported mutations via Sanger sequencing. The entire cohort of primary tumors and metastases was sequenced to detect the potential kinase domain mutations *PIK3C2B*^*G3542T*^ and *PIK3C2B*^*A3623G*^. Eventually, neither could be identified, or any other mutation in the conserved catalytic and activation loop motifs in exon 25 of *PIK3C2B* (Fig. 1D). Likewise, no alterations were found at amino acids positions 117 or 1469 (Fig. 1D). The only observed sequential deviations were already documented SNPs in exon 3 and exon 25 of *PIK3C2B* (Suppl. Fig. 1).

### PIK3C2B in silico

To put the results of the screening into a broader context, a set of 504 SQCC provided by the TCGA (cbioportal.org) was assessed in relation to *PIK3C2B* aberrations. As for somatic mutations, data were available for 177 tumors. Those harbored *PIK3C2B* alterations in 4 % (7/177) of all cases, which were non-redundant and spread across the gene. Interestingly, previously described alterations P117A and H1208R were also found in the cohort. *PIK3C2B* mutations were not associated with a poorer overall or disease-free survival prognosis (Fig. 2A). As for alterations in mRNA expression, a data set of 501 samples was available. Applying a z-score threshold of ± 1, the set was altered in 71/501 cases (upregulation in 50 cases, downregulation in 21 cases). Likewise, deviations in mRNA expression were not associated with significant deterioration of overall or disease-free survival (Fig. 2B). Also, there was no observable pattern between American Joint Committee on Cancer (AJCC) tumor stages and the appearance of somatic mutations or mRNA expression levels (Fig. 2C, n=328). Protein expression level measured by reverse phase protein arrays (RPPA) was not altered in any of the samples after applying a z-score of ± 1.

**Figure 2:**
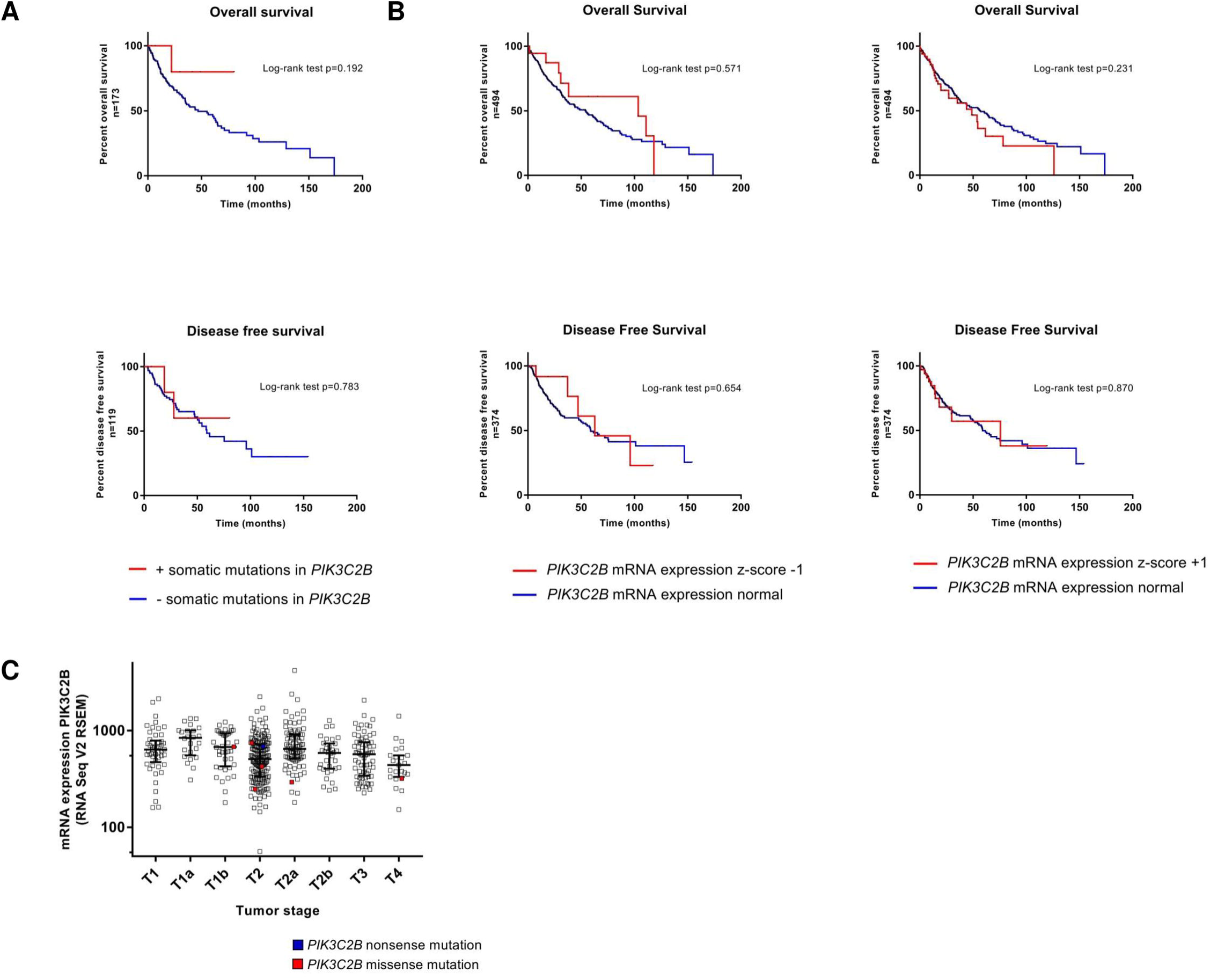
Meta-analysis of TCGA NSCLC-SQCC sequence data. **A** Kaplan-Meier estimates of overall and disease-free survival of patients with and without somatic mutations in *PIK3C2B*. **B** Kaplan-Meier estimates of overall and disease-free survival of patients with and without alterations in *PIK3C2B* mRNA expression (Z-score threshold ±1, RNA Seq V2 RSEM). **C** Scatter plot of *PIK3C2B* mRNA expression in all AJCC tumor stages (x-axis: AJCC tumor stages, y axis: RNA Seq V2 RSEM, log 10).

### Functional analysis of *PI3KC2β^C1181F^*/*PI3KC2β^H1208R^*

To assess the oncogenic potential of PI3KC2β mutations C1181F and H1208R, they were cloned into a Myc-tagged *PIK3C2B* expression vector via site directed mutagenesis. Successful sequence alteration was verified by Sanger sequencing (Fig. 3A). As a negative control, the ORF was removed to produce an empty backbone vector (EPIK2B). After CaCl_2_ transfection of HEK293 cells, *PIK3C2B* overexpression was examined on a transcriptional and translational level. Results of the conducted qPCRs and western blots indicated strong overexpression on both levels (Fig. 3B & 3C). Via immunofluorescence targeting the c-terminal Myc-tag of the protein, expression of exogenous PI3KC2β was witnessed in 65-75 % HEK cells after transfection (Fig. 3D).

**Figure 3:**
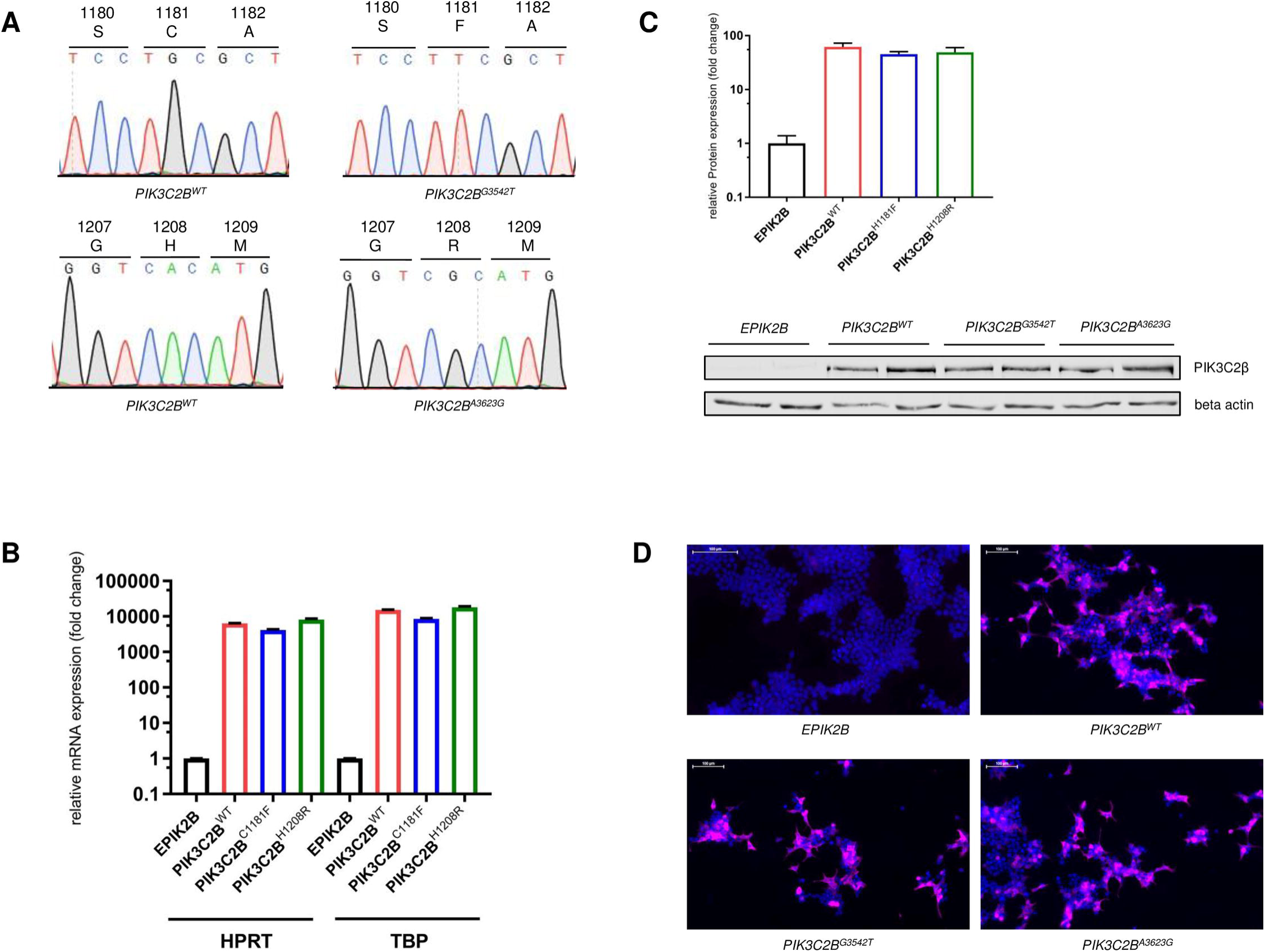
*PIK3C2B* plasmid engineering and validation. **A** Chromatograms of engineered *PIK3C2B* vectors. **B** Relative mRNA expression of *PIK3C2B* after transfection of HEK293 cells (48h) with plasmid constructs. **C** Relative protein expression of PI3KC2β after transfection of HEK293 cells (48h) with plasmid constructs. **D** Expression of exogenous PI3KC2β, visualized with immunofluorescence after transfection of HEK293 cells with plasmid constructs. Staining with DAPI (blue) and MycTag antibody (violet).

In addition, potential alterations in kinase activity caused by C1181F*/* H1208R were measured. Following immunoprecipitation of exogenously expressed PI3KC2β variants from HEK293 cells with a MycTag AB, kinase activity was measured with the ADP-Glo-Kinase Assay kit. Ultimately, no significant changes between PI3KC2β^WT^, PI3KC2β^C1181F^ and PI3KC2β^H1208R^ were detected (Fig. 4A).

**Figure 4:**
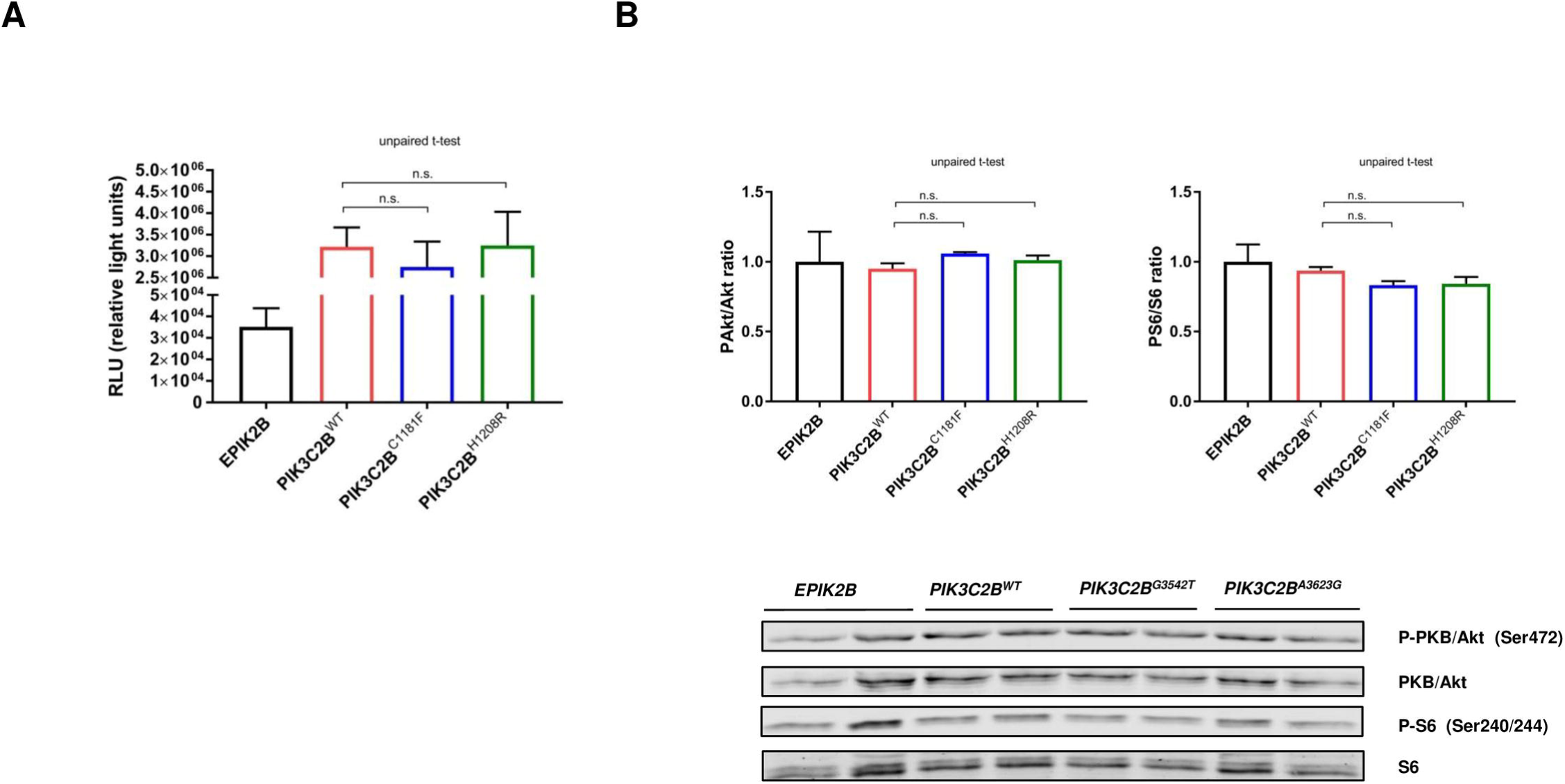
Functional Analysis of PI3KC2β^WT^, PI3KC2β^C1181F^ and PI3KC2β^H1208R^. **A** Lipid Kinase activity of PI3KC2β^WT^, PI3KC2β^C1181F^ and PI3KC2β^H1208R^ measured after immunoprecipitation of exogenously expressed PI3KC2β. **B** Ratios of Pakt/Akt and PS6/S6 after transfection of HEK293 cells with plasmid constructs. 12h after transfection, cells were starved for 24h with 0.5 % FCS.

Alterations in PI3K pathway activation were analyzed with western blots. HEK293 cells were transfected with plasmid variants and subsequently starved for 24 hours. Neither PI3KC2β^C1181F^ nor PI3KC2β^H1208R^ led to a significant increase in the phosphorylation of the pathway effector proteins PKB/Akt or S6 after transient overexpression compared to the wildtype allele (Fig. 4B).

## Discussion

Here, a large cohort of NSCLC-SQCC tumors was screened for four reported *PIK3C2B* missense mutations leading to amino acid exchanges (P117A, C1181F, H1208R, L1469F) (20). In addition, kinase domain mutations C1181F and H1208R were assessed for their oncogenic potential on a functional level.

The main focus lay on the two mutations in the kinase domain (C1181F, H1208R) bearing the highest potential for oncogenesis. None of these alterations were identified in 362 patient-derived tumor samples (Fig 1). Assuming the frequency found in Liu *et al*., a complete absence of the mutations by chance in the significantly larger cohort had a p-value < 0.0001. The screening was then extended to L1469F, a mutation described in the phosphoinositide-binding domain (PX-domain). After screening the first half of the cohort, no mutations were found in 163 samples. Finally, the cohort was also screened for P117A, a mutation that occurred in the proline-rich domain of the protein. This region has been proposed to play a role in kinase activity regulation and clathrin binding (29). There, the mutation was undetectable in 160 tested samples. The absence of any mutations in the cohort suggests that they might not confer a significant advantage in NSCLC-SQCC oncogenesis/signaling.

The lack of any detected mutations led to a thorough validation of the employed assay and the cohort. The technical approach was challenged by mixing different ratios of wildtype and mutant plasmids. Ultimately, the sequencing assay was found to be sensitive enough to detect the mutations at a 1:9 ratio (Fig. 1B). The obtained sensitivity was satisfying since the samples contained a minimum fraction of 30 % tumor tissue. Next, the cohort was challenged by screening it for p110_α_ mutations E542K and E545K. Both are charge reversing hotspot mutations that were proven to be oncogenic and frequently found in NSCLC-SQCC (25). There, both mutations were detected at the expected frequency (Fig 1C) according to the COSMIC database. These two experiments showed that the absence of detected mutations was neither due to a technical issue nor to a singularity of the cohort.

To put the findings of the screening into a broader context, a NSCLC-SQCC dataset provided by the TCGA was analyzed in regard to *PIK3C2B* alterations. As for somatic mutations (n=7/177), they were non-redundant, rare and did not change clinical outcomes (Fig.2A). Interestingly, P117A and H1208R were also present once in the cohort. One potential explanation could be that both are passenger mutations that are more frequent in the screened northern American cohorts. Alterations in mRNA expression (up or down) did likewise not change clinical outcomes (Fig.2B) and did not translate into overexpression at the protein level. Also, neither somatic mutations nor mRNA expression levels of *PIK3C2B* were associated with a particular tumor stage (Fig.2C).

In line with the findings of the screening, transient overexpression of the reported protein variants PI3KC2β^C1181F^ and PI3KC2β^H1208R^ in HEK293 cells (Fig. 3) did not reveal an effect of the mutations on lipid kinase activity (Fig. 4A) or PI3K pathway activation (Fig. 4B) in either direction. This shows that the two mutations do not confer any additional effect than the effect of the wildtype protein.

Taken together, the present data do not suggest a driver function of somatic *PIK3C2B* mutations in NSCLC-SCC and that aberrant PI3K pathway activation in NSCLC-SQCC occurs through alterations in more central compartments of the signaling axes like *EGFR, PIK3CA* and *PTEN* (25). Also, a recent study analyzed the mutational patterns in lung adenocarcinomas and squamous cell carcinomas (30). In accordance with the aforementioned findings, *PIK3C2B* was not found to be significantly mutated in either. As a more promising approach, future studies could investigate alterations in *PIK3C2B* concomitantly with alterations in other, potentially redundant PI3K isoforms.

In terms of cancer genetics, evidence for oncogenic implications of somatic *PIK3C2B* alterations is scarce. The only exception is a single nucleotide polymorphism that has been reported to be significantly associated with prostate cancer risk (31). So far, studies have mainly described amplifications of the genomic *PIK3C2B* locus. Gain at 1q32.1, the chromosomal region encoding for *PIK3C2B* and *MDM4* has been reported in studies assessing copy-number alterations in glioblastoma multiforme (32,33). In ovarian cancer, copy number gains of *PIK3C2B* have been reported as well (34).

Increased levels of cellular PI3KC2β have repeatedly been associated with oncogenesis. A study downregulating 779 kinases via RNAi in breast cancer cells (MCF7) ranked the siRNA targeting PI3KC2β as one of the top 20 to sensitize cells to tamoxifen (35). Another *in vitro* study overexpressing PI3KC2β in oesophageal squamous cells (Eca109) reached a similar conclusion. Overexpression of PI3KC2β led to a 4-fold reduction in sensitivity to cisplatin. siRNA mediated down-regulation of the enzyme resulted in restoration of sensitivity to the drug (17). Conversely though, promotion of resistance to thiopurines in leukemia cells through deletion of PI3KC2β has also been described (36). The effect of PI3KC2β may thus be drug specific.

*In vivo*, overexpression of PI3KC2β in suprabasal and basal epidermal cell layers in mice did not affect epidermal growth and differentiation (37). In the same study, mice with ubiquitous homozygous deletions of *PIK3C2B* were viable, fertile and without any reported phenotype.

Apart from cellular outcomes and phenotypes, the molecular consequences of PI3KC2β amplification on pathway signaling remain to be determined.

This task is complicated by the fact that PI3KC2β was discovered on the basis of sequence homologies rather than a functional context (38). Assuming that PI3KC2β is able to generate PtdIns(3,4)P_2_ (28), several studies investigated the possibility that the isoform is able to activate Akt kinase, a cardinal node in diverse signaling cascades. So far, contrasting evidence is present in the literature. Silencing of PI3KC2β has been shown to reduce Akt activation in neuroblastoma models (18). Conversely though, no effect on Akt phosphorylation was detected in epidermoid carcinoma cells (A-431) overexpressing PI3KC2β when compared to parental cells (16). Another study described an attenuation of Akt phosphorylation in PI3KC2β overexpressing HEK293 cells (39). To explain these seemingly contradictive effects on Akt activation, an indirect cross-talk mechanism with other signaling molecules not relying on the generation of specific phosphoinositides was proposed (40). Consistent with this hypothesis, a recent study found an indirect, even tissue specific effect of Akt activation upon PI3KC2β inhibition (41).

PI3KC2β may fulfill context dependent tasks in different cell types. Hence, it could pose a considerable challenge to determine direct downstream targets and the exact physiological conditions under which PI3KC2β acts. Nevertheless, it is a necessary prerequisite to integrate it into the precise context of cancer formation as it does not appear to be a classic oncogene.

## Acknowledgements

We would like to acknowledge the Microscopy Imaging Center of the University of Bern (MIC). Mr. Kind is enrolled in the Graduate School for Cellular and Biomedical Sciences (GCB) of the University of Bern that we would like to acknowledge for the training provided. Furthermore, we would like to acknowledge Fabiana Jacob and the entire staff of the DCR- VPH for a helping hand with the sequencing part. Also, the financial support of the Kinderkrebsstiftung Bern has to be highly acknowledged. Finally, the MT-Lab receives supports from the NCCR-TransCure.

---

**Suppl. Fig. 1:** **A** Incidence rate of detected SNP in *PIK3C2B.*

## Notes

**Disclosure of Potential Conflicts of Interest**: The authors have no conflict of interest regarding this publication.

## References

1. Castoria G, Migliaccio A, Bilancio A, Di Domenico M, de Falco A, Lombardi M, et al. PI3-kinase in concert with Src promotes the S-phase entry of oestradiol-stimulated MCF-7 cells. EMBO J. European Molecular Biology Organization; 2001;20:6050–9.

2. Sadhu C, Masinovsky B, Dick K, Sowell CG, Staunton DE. Essential Role of Phosphoinositide 3-Kinase δ in Neutrophil Directional Movement. J Immunol. 2003;170.

3. Graupera M, Guillermet-Guibert J, Foukas LC, Phng L-K, Cain RJ, Salpekar A, et al. Angiogenesis selectively requires the p110α isoform of PI3K to control endothelial cell migration. Nature. Nature Publishing Group; 2008;453:662–6.

4. Datta SR, Dudek H, Tao X, Masters S, Fu H, Gotoh Y, et al. Akt phosphorylation of BAD couples survival signals to the cell-intrinsic death machinery. Cell. 1997;91:231–41.

5. Makarov SS, Romashkova JA. NF-kappaB is a target of AKT in anti-apoptotic PDGF signalling. Nature. 1999;401:86–90.

6. Fruman DA, Rommel C. PI3K and cancer: lessons, challenges and opportunities. Nat Rev Drug Discov [Internet]. 2014;13:140–56.

7. Stambolic V, Suzuki A, de la Pompa JL, Brothers GM, Mirtsos C, Sasaki T, et al. Negative regulation of PKB/Akt-dependent cell survival by the tumor suppressor PTEN. Cell. 1998;95:29–39.

8. Li J, Yen C, Liaw D, Podsypanina K, Bose S, Wang SI, et al. PTEN, a putative protein tyrosine phosphatase gene mutated in human brain, breast, and prostate cancer. Science (80-) [Internet]. 1997/03/28. 1997;275:1943–7.

9. Samuels Y, Ericson K. Oncogenic PI3K and its role in cancer. Curr Opin Oncol. 2006;18:77–82.

10. Samuels Y. High frequency of mutations of the PIK3CA gene in human cancers. Science (80-). 2004;304:554.

11. Kang S, Bader AG, Vogt PK. Phosphatidylinositol 3-kinase mutations identified in human cancer are oncogenic. Proc Natl Acad Sci. 2005;102:802–7.

12. Zhao JJ, Liu Z, Wang L, Shin E, Loda MF, Roberts TM. The oncogenic properties of mutant p110 and p110 phosphatidylinositol 3-kinases in human mammary epithelial cells. Proc Natl Acad Sci. 2005;102:18443–8.

13. Piddock RE, Loughran N, Marlein CR, Robinson SD, Edwards DR, Yu S, et al. PI3Kδ and PI3Kγ isoforms have distinct functions in regulating pro-tumoural signalling in the multiple myeloma microenvironment. Blood Cancer J. 2017;7:e539.

14. Ikeda H, Hideshima T, Fulciniti M, Perrone G, Miura N, Yasui H, et al. PI3K/p110{delta} is a novel therapeutic target in multiple myeloma. Blood. American Society of Hematology; 2010;116:1460–8.

15. Yoshioka K, Yoshida K, Cui H, Wakayama T, Takuwa N, Okamoto Y, et al. Endothelial PI3K-C2α, a class II PI3K, has an essential role in angiogenesis and vascular barrier function. Nat Med. 2012;18:1560–9.

16. Katso RM, Pardo OE, Palamidessi A, Franz CM, Marinov M, De Laurentiis A, et al. Phosphoinositide 3-Kinase C2beta Regulates Cytoskeletal Organization and Cell Migration via Rac-dependent Mechanisms. Mol Biol Cell. 2006;17:3729–44.

17. Liu Z, Sun C, Zhang Y, Ji Z, Yang G. Phosphatidylinositol 3-Kinase-C2β Inhibits Cisplatin-Mediated Apoptosis via the Akt Pathway in Oesophageal Squamous Cell Carcinoma. J Int Med Res. 2011;39:1319–32.

18. Russo A, Okur MN, Bosland M, O’Bryan JP. Phosphatidylinositol 3-kinase, class 2 beta (PI3KC2β) isoform contributes to neuroblastoma tumorigenesis. Cancer Lett. 2015;359:262–8.

19. Boller D, Doepfner KT, De Laurentiis A, Guerreiro AS, Marinov M, Shalaby T, et al. Targeting PI3KC2β impairs proliferation and survival in acute leukemia, brain tumours and neuroendocrine tumours. Anticancer Res. 2012;32:3015–27.

20. Liu P, Morrison C, Wang L, Xiong D, Vedell P, Cui P, et al. Identification of somatic mutations in non-small cell lung carcinomas using whole-exome sequencing. Carcinogenesis. 2012;33:1270–6.

21. Pesch B, Kendzia B, Gustavsson P, Jöckel K-H, Johnen G, Pohlabeln H, et al. Cigarette smoking and lung cancer-relative risk estimates for the major histological types from a pooled analysis of case-control studies. Int J Cancer. Wiley Subscription Services, Inc., A Wiley Company; 2012;131:1210–9.

22. Bass AJ, Watanabe H, Mermel CH, Yu S, Perner S, Verhaak RG, et al. SOX2 is an amplified lineage-survival oncogene in lung and esophageal squamous cell carcinomas. Nat Genet. 2009;41:1238–42.

23. Singh A, Misra V, Thimmulappa RK, Lee H, Ames S, Hoque MO, et al. Dysfunctional KEAP1-NRF2 interaction in non-small-cell lung cancer. Meyerson M, editor. PLoS Med. 2006;3:e420.

24. Rekhtman N, Paik PK, Arcila ME, Tafe LJ, Oxnard GR, Moreira AL, et al. Clarifying the Spectrum of Driver Oncogene Mutations in Biomarker-Verified Squamous Carcinoma of Lung: Lack of EGFR/KRAS and Presence of PIK3CA/AKT1 Mutations. Clin Cancer Res. 2012;18:1167–76.

25. Hammerman PS, Lawrence MS, Voet D, Jing R, Cibulskis K, Sivachenko A, et al. Comprehensive genomic characterization of squamous cell lung cancers. Nature. 2012;489:519–25.

26. Cerami E, Gao J, Dogrusoz U, Gross BE, Sumer SO, Aksoy BA, et al. The cBio Cancer Genomics Portal: An open platform for exploring multidimensional cancer genomics data. Cancer Discov. 2012;2:401–4.

27. Gao J, Aksoy BA, Dogrusoz U, Dresdner G, Gross B, Sumer SO, et al. Integrative Analysis of Complex Cancer Genomics and Clinical Profiles Using the cBioPortal. Sci Signal. 2013;6:pl1-pl1.

28. Arcaro A, Volinia S, Zvelebil MJ, Stein R, Watton SJ, Layton MJ, et al. Human phosphoinositide 3-kinase C2beta, the role of calcium and the C2 domain in enzyme activity. J Biol Chem. 1998;273:33082–90.

29. Wheeler M, Domin J. The N-terminus of phosphoinositide 3-kinase-C2? regulates lipid kinase activity and binding to clathrin. J Cell Physiol. 2006;206:586–93.

30. Campbell JD, Alexandrov A, Kim J, Wala J, Berger AH, Pedamallu CS, et al. Distinct patterns of somatic genome alterations in lung adenocarcinomas and squamous cell carcinomas. Nat Genet. 2016;48:607–16.

31. Koutros S, Schumacher FR, Hayes RB, Ma J, Huang W-Y, Albanes D, et al. Pooled Analysis of Phosphatidylinositol 3-Kinase Pathway Variants and Risk of Prostate Cancer. Cancer Res. 2010;70:2389–96.

32. Sumihito N, Joel L, Anne W, Young Ho K, Jian H, Catherine L, et al. Intratumoral patterns of genomic imbalance in glioblastoma. Brain Pathol. 2010;20:936–44.

33. Rao SK, Edwards J, Joshi AD, Siu I-M, Riggins GJ. A survey of glioblastoma genomic amplifications and deletions. J Neurooncol. 2010;96:169–79.

34. Zhang L, Huang J, Yang N, Greshock J, Liang S, Hasegawa K, et al. Integrative Genomic Analysis of Phosphatidylinositol 3'-Kinase Family Identifies PIK3R3 as a Potential Therapeutic Target in Epithelial Ovarian Cancer. Clin Cancer Res. 2007;13:5314–21.

35. Iorns E, Lord CJ, Ashworth A. Parallel RNAi and compound screens identify the PDK1 pathway as a target for tamoxifen sensitization. Biochem J. 2009;417:361–71.

36. Diouf B, Cheng Q, Krynetskaia NF, Yang W, Cheok M, Pei D, et al. Somatic deletions of genes regulating MSH2 protein stability cause DNA mismatch repair deficiency and drug resistance in human leukemia cells. Nat Med. 2011;17:1298–303.

37. Harada K, Truong AB, Cai T, Khavari PA. The class II phosphoinositide 3-kinase C2beta is not essential for epidermal differentiation. Mol Cell Biol. American Society for Microbiology (ASM); 2005;25:11122–30.

38. Brown RA, Ho LKF, Weber-Hall SJ, Shipley JM, Fry MJ. Identification and cDNA Cloning of a Novel Mammalian C2 Domain-Containing Phosphoinositide 3-Kinase, HsC2-PI3K. Biochem Biophys Res Commun. 1997;233:537–44.

39. Domin J, Harper L, Aubyn D, Wheeler M, Florey O, Haskard D, et al. The class II phosphoinositide 3-kinase PI3K-C2beta regulates cell migration by a PtdIns3P dependent mechanism. J Cell Physiol. 2005;205:452–62.

40. Falasca M, Maffucci T. Regulation and cellular functions of class II phosphoinositide 3-kinases. Biochem J. 2012;443:587–601.

41. Alliouachene S, Bilanges B, Chicanne G, Anderson KE, Pearce W, Ali K, et al. Inactivation of the Class II PI3K-C2β Potentiates Insulin Signaling and Sensitivity. Cell Rep. Elsevier; 2015;13:1881–94.

